# “The Brain is…”: A Survey of The Brain’s Many Definitions

**DOI:** 10.1101/2023.10.26.564205

**Authors:** Taylor Bolt, Lucina Q. Uddin

## Abstract

A reader of the peer-reviewed neuroscience literature will often encounter expressions like the following: ‘the brain is a dynamic system’, ‘the brain is a complex network’, or ‘the brain is a highly metabolic organ’. These expressions attempt to define the essential functions and properties of the mammalian or human brain in a simple phrase or sentence, sometimes using metaphors. We sought to survey the most common phrases of the form ‘the brain is…’ in the biomedical literature to provide insights into current conceptualizations of the brain. Utilizing text analytic tools applied to a large sample (> 4 million) of peer-reviewed full-text articles and abstracts, we extracted several thousand phrases of the form ‘the brain is…’ and identified over a dozen frequently appearing phrases. The most used phrases included metaphors (e.g., the brain as a ‘information processor’ or ‘prediction machine’) and descriptions of essential functions (e.g., ‘a central organ of stress adaptation’) or properties (e.g., ‘a highly vascularized organ’). Comparison of these phrases with those involving other bodily organs (e.g. the heart, liver, etc.) highlighted common phrases between the brain and other organs, such as the heart as a ‘complex dynamic system’. However, the brain was unique among organs in the number and diversity of metaphors ascribed to it. The results of our analysis underscore the diversity of qualities and functions commonly attributed to the brain in the biomedical literature and suggest a range of conceptualizations that defy unification.

## Introduction

The heart is a pump sending blood around the body, the primary organ of the circulatory system, and a muscular organ made up of cardiac muscle. While these definitions invariably fail to appreciate the complexity of the heart’s operations, they provide simple answers to a simple question: what sort of thing is a heart? They function as pedagogical or explanatory tools that convey the heart’s ‘essential’ functions and qualities. Some of these descriptions express analogies (e.g., ‘the heart is a pump’), others express part-whole relationships (e.g., the ‘primary organ of the circulatory system’), while others highlight the essential biochemical or cellular properties of the organ that enable its function (e.g., ‘made up of cardiac muscle’).

Not all organs of the human body can be so readily defined. Communicating the essential functions and qualities of the human or mammalian brain is a notoriously difficult task. For many scientists and philosophers, the brain’s essential qualities and functions are a matter of substantive scientific interest. At stake is not merely linguistic disagreements, but the guiding interpretive framework for communicating neuroscientific results and potentially the favored methodologies by which we collect relevant data (Kelty-Stephen et al., 2022). For example, the computational or information-processing analogy has inspired a wealth of research findings and interpretive frameworks in cognitive, systems and computational neuroscience (Cobb, 2020). Favored definitions may be non-metaphorical as well, inspired by the brain’s primary role in a physiological process (e.g., the brain’s role in hormone regulation and stress).

To appreciate the diversity of definitions that scientists have ascribed to the brain in attempts to study it and determine whether unifying conceptualizations are evident, we conducted a survey of the biomedical literature (PubMed Central Open Access) using natural language processing (NLP) techniques. We searched for noun phrases following variations of the phrase ‘The brain is a…’. Our survey targeted expressions where the subject (‘the brain’) is linked via a copular verb (‘is’) to a predicative expression consisting of a determiner (‘the/a’) followed by a noun phrase (e.g. ‘complex computer’). An example expression that would fit this pattern is ‘the brain is a complex computer’. NLP-based semantic embeddings and dimension-reduction techniques identified over a dozen commonly used copular expressions to describe the brain, many with quite different meanings. Some of these expressions were metaphors (‘the brain is a computer’), others described an essential property (‘the brain is a metabolically expensive organ’), while others described an essential function (‘the brain is the key regulator of stress’). Comparison with expressions involving other bodily organs showed that some expressions were not unique to the brain. Our results underscore the diversity of attributes that scientists have ascribed to the brain in the biomedical literature.

## Results

The biomedical corpus for analysis consisted of full text articles from the Pubmed Central Open Access Subset (N=4,993,411) and abstracts from leading neuroscience journals (N=253,022). Of the total corpus, 4,375 expressions from 897 peer-reviewed journals were found that matched expressions of the form ‘the brain is …’. Due to the disproportionate representation of large open-access journals in the corpus, journals such as *PLoS ONE* (N = 208), *International Journal of Molecular Sciences* (N = 187), *Frontiers in Neuroscience (*N *=* 172), *Psychology (*N = 130), *Human Neuroscience (*N = 105), and *Scientific Reports* (N = 70) constituted the largest share of matched expressions. Articles from the current decade (>= 2020) constituted the largest share of matched expressions (∼56%). Of the 2,204 (51%) matched expressions identified from sections of text with ‘standard’ titles (e.g., titles containing ‘Abstract’, ‘Introduction’, ‘Results’, ‘Discussion’, ‘Methods’, etc.), 54% were found in introduction sections, 22% were found in discussions, and 16% were found in abstracts.

Following extraction, the text in each matched expression was converted to a vector space via a pretrained embedding model (Deka et al., 2022). To identify commonly used expressions, we applied dimensionality reduction to the embedding space to obtain two dimensions using UMAP (McInnes et al., 2020) and clustered the phrases using a hierarchical density-based clustering algorithm (HDBSCAN; (Campello et al., 2013) (**Figure 1**). Of the 26 clusters extracted by the HDBSCAN algorithm (see *Methods and Materials*), 21 were found to constitute semantically coherent groups of expressions – i.e., groups of expressions that express similar meaning. Two clusters (18 & 19) were found to contain similar meanings and were merged, leaving a total of 20 semantically coherent clusters. Labels for each cluster were generated from manual inspection. In terms of overall organization of the semantic embedding space, the phrases were organized along a single dominant dimension, such that abstract/metaphorical phrases were concentrated in the top-left, to more cellular/biochemical phrases in the bottom-right.

**Figure 1.**
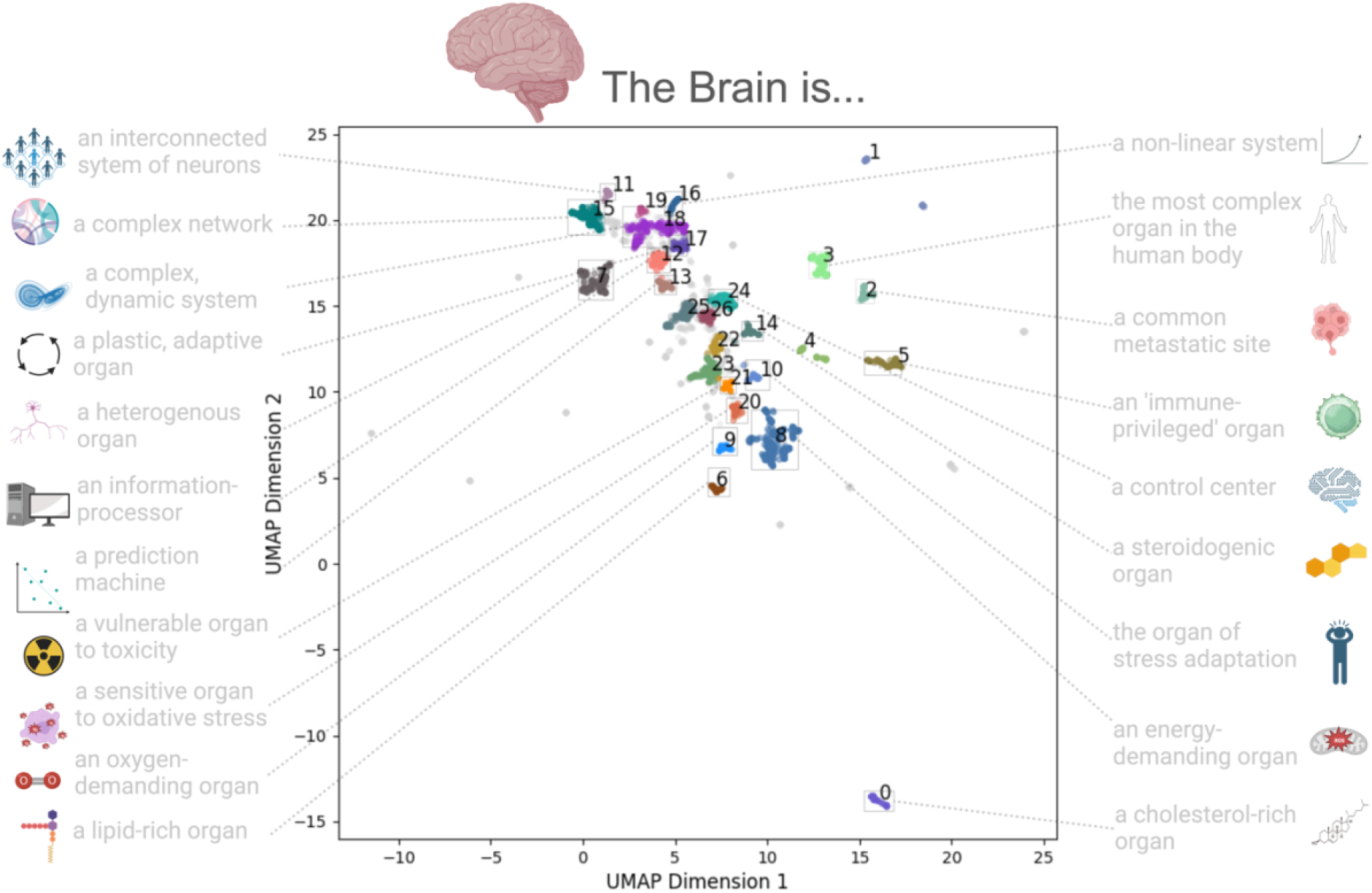
‘The brain is…’ Expressions in a Two-Dimensional Embedding Space. Expressions matching the form ‘The brain is…’ embedded into a two-dimensional space via a dimension-reduction (UMAP) applied to their semantic embeddings. The distance between points in this space reflect the semantic similarity between the expressions – i.e., expressions (points) in this space that are closer together reflect similar meanings. Expressions are color-coded according to their cluster assignment from the HDBSCAN clustering algorithm. Each semantically coherent cluster (N=21) is labeled by a manual interpretation of the expressions in the cluster.

Commonly used noun phrases following the phrase ‘the brain is…’ span multiple levels of organization, from the biochemical (e.g., ‘a lipid rich organ’) to the structural (e.g., ‘a heterogenous organ’); as well as different types of expressions, including metaphors (e.g., ‘a prediction machine’ and ‘a control system’). Of the 20 semantically coherent clusters, the top clusters were (in descending order), the brain is ‘an energy demanding organ’ (N = 461), ‘a complex, dynamic system’ (N = 301), ‘a complex network’ (N = 277), ‘a heterogenous organ (N = 257), and ‘a control system’ (N = 160).

The occurrence of each expression was unequally distributed across journals **(Supplementary Table 1**), with expressions clustering into journals with distinct disciplinary concentrations. Some of these were predictable – e.g., biochemical expressions, such as ‘cholesterol-’ and ‘lipid-rich’ tended to appear more frequently in journals with a focus in biochemistry and molecular biology (e.g. *Oxidative Medicine and Cellular Longevity, Molecules*), and mentions of the brain as a ‘common site of metastasis’ appear more frequently in oncology journals (e.g. *Cancers, Frontiers in Oncology*).

Expressions of metaphors tended to appear more frequently in psychology and human neuroscience journals. For example, most of the expressions found in the journal *Frontiers in Psychology* were the brain as a ‘prediction machine’ or ‘computer/information processor’. The expressions of the brain as a ‘non-linear’ or ‘complex, dynamic’ system and ‘complex network’ tended to occur in human neuroscience/neuroimaging journals, including *Human Brain Mapping, Frontiers in Human Neuroscience, Neuroimage, Brain and Behavior*, and *Network Neuroscience*.

A further question is whether the expressions attributed to the brain are unique to that organ or appear in reference to other bodily organs. Using the same methodology, we separately embedded and clustered expressions where the subject was the lungs, kidneys, pancreas, heart, stomach, or liver. Interestingly, we found that all organs surveyed, except for the stomach, contained expressions of the form the organ is a ‘complex’ or ‘dynamic’ organ. For example, the heart is variously referred to as both a ‘complex’ and a ‘dynamic’ organ in the biomedical literature (**Supplementary Figure 1**). In addition, expressions involving the frequency (or infrequent) occurrence of metastases were found across all organs. Other overlapping expressions between the brain and other organs, included the kidneys and heart as an ‘energy-demanding organ’ and the liver as a ‘heterogenous’ and ‘immune-privileged’ organ. However, the number and diversity of metaphorical expressions involving the brain was distinct among all organs surveyed. For example, metaphors such as the brain is a ‘prediction machine’, ‘information processor’, and a ‘control system’ were unique to the brain.

## Discussion

What is the brain? The results of our analysis underscore the diversity of qualities and functions commonly attributed to the human or mammalian brain in the biomedical literature. For scientists in some disciplines, the brain is understood from an abstract level, in the form of analogy – a computer/information processor, a prediction (Bayesian) machine, or a dynamic system. For others, the brain is a site of remarkable functional capacities, including its role in stress adaptation, its status as a relatively ‘immune-privileged’ organ, and its high metabolic activity. For others still, the brain is defined in terms its biochemical properties as a lipid-and cholesterol-rich organ or its vulnerability to oxidative stress.

The brain is not entirely unique in the predicates attributed to it. The body is filled with other ‘complex’ and ‘dynamic’ organs, according to the biomedical literature. These observations remind us that the ‘complex system’ that constitutes the brain is embedded in a much larger complex system that constitutes the human body (Varela et al., 1992). Bodily organs, such as the heart, kidneys, lungs and liver, are also complex systems that in some respects, rival that of the brain. Any satisfactory conceptualization of the brain’s function and properties should consider the interconnections between the brain and these organs. However, we found that the number and diversity of metaphors ascribed to the brain was much greater in comparison to other bodily organs. This observation underscores the fact that the brain’s function(s) lacks a unifying conceptualization compared with other bodily organs.

The appearance of this diversity of expression in the biomedical literature raises the question of implications for scientific communication. In many cases, it’s clear that these varied expressions do serve a purpose: expressions such as the ‘the brain is a highly adaptive, plastic organ’ or ‘the brain is an oxygen-demanding organ’ communicate an attribute of the brain that enables its unique functions or demarcates it from other organs in the body. Others attempt to communicate the essential unique function or property of the brain that makes it what it is – e.g., ‘the brain is an information processor/computer’ or ‘the brain is a dynamic, non-linear system’. In many cases, these analogical expressions function to justify the author’s research approach or perspective, sometimes in contrast to a rival approach or perspective.

Importantly, these expressions often serve to introduce a research or review article, as evidenced by the fact they often appear in the Introduction section of articles. They are not propositions to be assessed by empirical observations or experiments reported in the research article. Rather, they serve to introduce more concrete and testable hypotheses, detailed frameworks and theories, and/or methodological approaches.

We agree with and reiterate a common expression in the biomedical literature: the brain is the most complex organ of the human body. This complexity is not only found in its biological structure, but also in the predicates that scientists use to describe it. The fact that a range of conceptualizations exist that defy unification has implications for science communication, neuroscience education, and society more broadly.

### Supplementary Materials

**Supplementary Figure 1.**
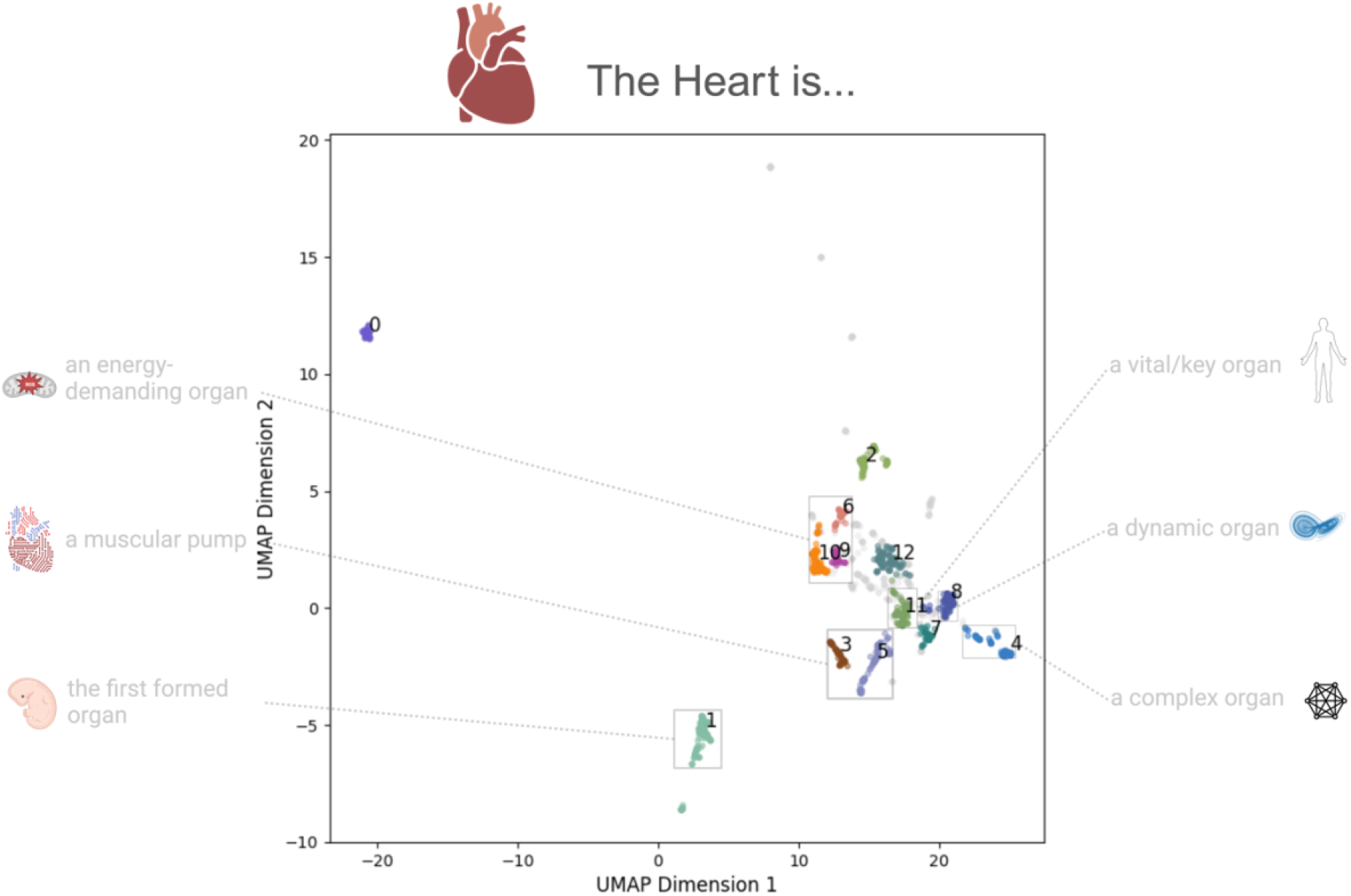
‘The heart is…’ Expressions in a Two-Dimensional Embedding Space. Expressions matching the form ‘The heart is…’ embedded into a two-dimensional space via a dimension-reduction (UMAP) applied to their semantic embeddings. The distance between points in this space reflect the semantic similarity between the expressions – i.e., expressions (points) in this space that are closer together reflect similar meanings. Expressions are color-coded according to their cluster assignment from the HDBSCAN clustering algorithm. Each semantically coherent cluster (N=9) is labeled by a manual interpretation of the expressions in the cluster. Boxes including multiple clusters are mapped to the same label.

**Supplementary Table 1.**
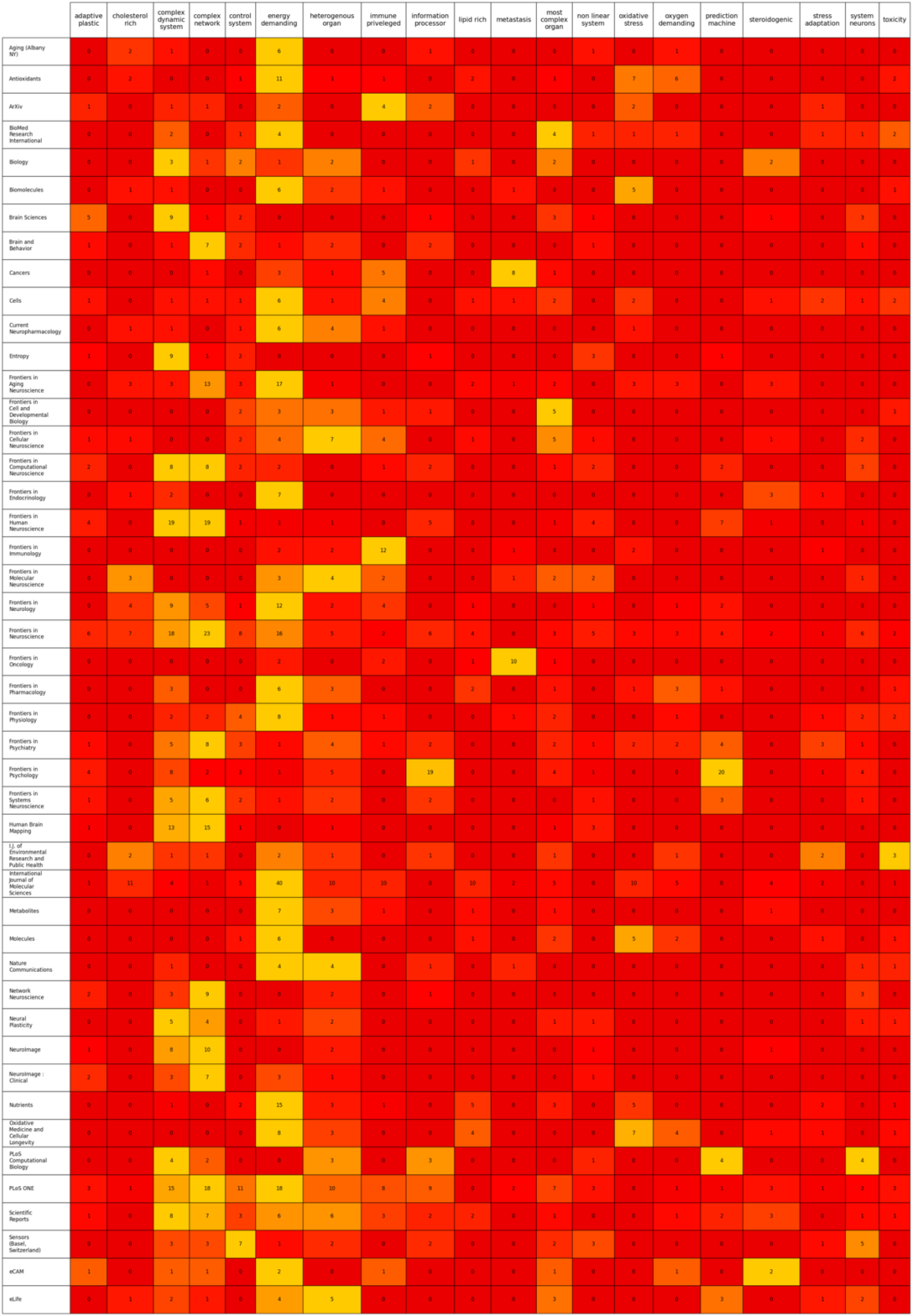
Counts of Expressions from Each Cluster by Journal Title. A cross-tabulation table containing the counts of expressions in each cluster (column) by journal title (row). Only journals containing >= 20 expressions were included in the table. Clusters are labeled with the same labels as appears in **Figure 1** (abbreviated for space). Cells of the table are color-coded according to their row-wise normalized (min-max normalization) counts, with lighter colors corresponding to more frequent counts relative to the total counts in each journal (row).

**Supplementary Table 2.** Neuroscience Journal Abstracts Included in Corpus. The count of abstracts of neuroscience journals included in corpus for analysis.

| Journal | Abstract Count |
| --- | --- |
| Journal of Neuroscience | 37967 |
| Journal of Cognitive Neuroscience | 4227 |
| Neuroscience Letters | 35108 |
| Neuropsychologia | 8996 |
| Neuroimage | 22060 |
| Nature Reviews Neuroscience | 1319 |
| Journal of Neurophysiology | 19269 |
| Trends in Cognitive Science | 2316 |
| Neuroscience and Biobehavioral Reviews | 5492 |
| Human Brain Mapping | 5911 |
| Psychophysiology | 4178 |
| Cortex | 4875 |
| Experimental Brain Research | 14804 |
| Nature Reviews Neurology | 888 |
| European Journal of Neuroscience | 13157 |
| Neuron | 11945 |
| The Neuroscientist | 1083 |
| Annual Review of Neuroscience | 696 |
| Molecular Neurobiology | 5933 |
| Brain | 8470 |
| Social Cognitive and Affective Neuroscience | 1980 |
| Nature Neuroscience | 5054 |
| Journal of Neurology | 9330 |
| Behavioral and Brain Sciences | 3465 |
| Progress in Neurobiology | 1921 |
| Biological Psychiatry | 9314 |
| Trends in Neuroscience | 3003 |
| Current Opinion in Neurology | 2644 |
| Cerebral Cortex | 7617 |

## Methods and Materials

### Dataset

The corpus used for analysis consisted of all full-text articles from the PubMed Central (PMC) Open Access subset (https://www.ncbi.nlm.nih.gov/pmc/tools/openftlist/) and the PMC Author Manuscript Dataset as of July 23, 2023 (N=4,993,411). The PMC Open Access Subset provides access to full texts from open access peer-reviewed journals. The PMC Author Manuscript Dataset provides access to full texts of manuscripts made available in PMC by authors in compliance with the NIH Public Access Policy. Both sources form part of PMC’s Open Access Collection (https://www.ncbi.nlm.nih.gov/pmc/tools/textmining/). Bulk downloads of the full Open Access Collection articles were conducted using the PMC FTP service. To supplement our corpus with scientific text outside the PMC Open Access Collection, we pulled abstracts from the PubMed database from select neuroscience journals (N_journals_ = 29, N_articles_ = 253,022; **Supplementary Table 2**) as of August 26, 2023 via the EFetch utility (Sayers, 2022). Overall, a total of approximately 5 million articles were downloaded and screened for our analysis. All code for preprocessing and analysis are provided at https://github.com/tsb46/the_brain_is.

### Preprocessing of PMC Open Access Collection

Full-text articles from the PMC Open Access Collection were downloaded in XML format. To parse the article XML files into structured data for analysis we used the Pubmed Parser package in Python (Achakulvisut et al., 2020). As an initial filtering step to reduce the number of articles in the next stage of preprocessing, the text of each section in the XML article files were searched for mentions of the entity ‘brain’, ‘lung(s)’, ‘pancreas’, ‘stomach’, ‘liver’, ‘kidney(s)’ or ‘heart’ (case insensitive and lemmatized) via the entity extraction pipeline in ScispaCy (Neumann et al., 2019; *en_core_sci_sm* model; v0.5.2). This initial filtering step reduced the size of the PMC Open Access Collection to 1,488,212 articles.

### Detection of Copular Expressions of the Form ‘The organ is…’

We first sought to find sentences containing phrases/expressions of the form ‘the organ is…’ in our corpus, where the organ label is ‘brain’, ‘lung(s)’, ‘pancreas’, ‘stomach’, ‘liver’, ‘kidney(s)’ or ‘heart’. We used the *PhraseMatcher* utility in spaCy package (https://github.com/explosion/spaCy) to detect sequences of tokens in text with prespecified linguistic properties, such as string matching, part-of-speech tags, dependency tree labels. The phrase we sought to detect in the corpus has a specific linguistic structure: the organ token (e.g. ‘brain’, ‘heart’) as the nominal subject of the clause (case insensitive match) preceded by a determiner (the/a) and followed by the copular verb (‘is’). In addition, this phrase (e.g. ‘The organ is…’) is followed by another determiner (the/a) that precedes the noun phrase. Hypothetical phrases that match this pattern would be ‘the brain is a complex system’ or ‘the heart is a muscular pump’. To capture adjectives that modify or further describe the noun, such as the ‘human brain’ or ‘mammalian brain’, we also allowed for matches with an adjective token preceding the token ‘brain’. Tokenization, dependency parsing and part-of-speech tagging were performed with the *en_core_sci_lg* model in ScispaCy (v0.5.2; Neumann et al., 2019). The *PhraseMatcher* pipeline applied to the filtered PMC Open Access Collection articles (N = 561,836) and neuroscience journal abstracts (N=253,022) identified 25,800 sentences containing a sequence of tokens that matched the token sequence template described above.

### Information Extraction of Matched Expressions

The *PhraseMatcher* pipeline detected the noun phrase and copular verb (‘is’) that forms the subject of the sentence, the phrase ‘The organ is…’. To extract the entire expression, including the full noun phrase(s) that follow the phrase ‘The organ is…’, further text parsing was performed. For accurate quantification of semantic similarity, it was important to extract the relevant tokens of the noun phrases following the phrase ‘The organ is…’, or irrelevant tokens/words would contribute to the semantic similarity estimates. One possibility is to extract the entire sentence containing the matched expressions, but this was found to include too much irrelevant information. For example, extracting the entire sentence: ‘The brain is a complex system, and this is the inspiration for our analysis in the manuscript’ would include the irrelevant second clause (‘and this is the inspiration…’). We developed a custom rule-based algorithm based on traversal of the dependency tree extracted from the sentence by the ScispaCy model (https://spacy.io/api/dependencyparser). Briefly, the algorithm consisted of the following primary steps: 1) start with the immediate rightward children tokens of the copular verb ‘is’, 2) loop through the children tokens and detect any instance of a nominal modifier, conjunction, or relative clause modifier (in that order) syntactic relationship between the head and the child token. 3) If one token with any of these syntactic relationships is found, extract the immediate rightward children of that token, and repeat the process. Minor modifications of this algorithm were included based on repeated trial-and-error experiments on the matched expressions. The full number of matched expressions extracted from this algorithm was 24,427. The number of matched expressions that included the ‘brain’ as the subject was 4,357. The full list of the 24,427 expressions extracted from this algorithm are available at (https://github.com/tsb46/the_brain_is/blob/main/data/matched_expressions.txt).

### Phrase Embedding and Clustering

The primary goal of our analysis was to estimate the common types of ‘The brain is…’ expressions in the biomedical literature. To identify different types of expressions, we grouped the 4,836 expressions matching ‘the brain is…’ into groups with identical or very similar semantic meaning using a clustering approach. First, the expressions (extracted from the information extraction pipeline described above) were transformed into a semantic embedding space (N_dimensions_ = 768) via a pretrained neural network model, previously trained on semantic similarity tasks for scientific text (Deka et al., 2022; https://huggingface.co/pritamdeka/S-Scibert-snli-multinli-stsb). The expressions in the embedding vector space were then fed to the uniform manifold approximation algorithm (UMAP; McInnes et al., 2020) for dimension reduction to two dimensions. Two central parameters – the number of nearest neighbors in initial graph construction and the minimum distance points are allowed to be apart -control the coordinates and global structure of the semantic expressions in the dimension-reduced space. Based on visualization of all expressions in the two-dimensional space, the following parameters were found to produce Euclidean distances between expressions that most closely followed the semantic similarity/dissimilarity in the expressions (*N*_*Neighbors*_ = 15; *min*_*dist*_ = 0).

The two-dimensional coordinates of each expression in the dimension-reduced space were then fed to a hierarchical density-based clustering algorithm (HDBSCAN; McInnes et al., 2017) to isolate groups of expressions with very similar or identical semantic meanings. The number of clusters extracted from the HDBSCAN algorithm is controlled indirectly by setting the minimum cluster size. Manual inspection of cluster solutions across varying values of this parameter revealed that a minimum cluster size of 50 maximized semantic interpretability.

For comparison with ‘brain’ expressions, expressions containing other organs (e.g. heart, kidneys) as the subject were run through the same algorithm. The same UMAP parameters were used, but the minimum cluster size was adjusted to each set of expressions for maximized semantic interpretability. Note, excluding ‘heart’ expressions (**Supplementary Figure 1**), the UMAP visualizations (**Figure 1**) for other organs were not included in the manuscript, but UMAP results/plots (and parameters) for each organ can be found at (https://github.com/tsb46/the_brain_is/blob/main/the_brain_is.ipynb).

